# Comparative analysis of plasma and bone marrow nutrient levels in pediatric B-ALL patients

**DOI:** 10.1101/2025.08.05.668707

**Authors:** Keene L. Abbott, Ahmed Ali, Muhammad Bin Munim, Devika P. Bagchi, Tenzin Kunchok, Millenia Waite, Caroline R. M. Wiggers, Boris Dimitrov, Marian H. Harris, Birgit Knoechel, Lewis B. Silverman, Matthew G. Vander Heiden, Yana Pikman

## Abstract

Nutrient availability in the tumor microenvironment is a key determinant of cancer progression and therapeutic response, yet the physiological nutrient environment for most cancers is poorly understood. In this study, we investigated nutrient levels in pediatric B-cell acute lymphoblastic leukemia (B-ALL) patients across different subtypes undergoing chemotherapy, focusing on both bone marrow and circulation. Our analysis revealed distinct differences in nutrient profiles between leukemic and healthy plasma and among B-ALL subtypes, with hyperdiploid B-ALL exhibiting pronounced alterations in arginine and asymmetric dimethylarginine metabolism. Bone marrow and blood plasma exhibited largely similar metabolite profiles, even after chemotherapy, indicating these environments are metabolically comparable. Comparisons with renal cell carcinoma and non-small cell lung cancer highlighted a unique enrichment of tricarboxylic acid cycle intermediates in the circulation of B-ALL patients. These findings provide a comprehensive view of nutrient dynamics in pediatric B-ALL and identify metabolic alterations that could guide biomarker discovery and new therapeutic strategies.

## Introduction

Acute lymphoblastic leukemia (ALL) remains the most frequently diagnosed pediatric malignancy, comprising 22% of cancers in children under 15 years of age^1,2^. The current treatment of childhood ALL represents a remarkable success story in modern medicine, with survival rates rising from 10% in the 1960s to more than 90% today^1,3^. This progress can be attributed to effective risk stratification at diagnosis, improved multi-agent chemotherapy regimens, measurement and treatment of minimal residual disease, and better supportive care.

Although deeper understanding of cell-intrinsic factors like genetic mutations has resulted in improved therapeutic outcomes, certain high-risk subtypes of ALL, including those with hypodiploidy, KMT2A rearrangements or Philadelphia chromosome (Ph) positivity, remain therapeutically challenging with relatively poor prognoses despite intensification of treatment regimens or addition of targeted therapies^1^. Further, although overall survival rates for B-ALL have dramatically improved in the last fifty years, limited progress has been made in outcomes for the 10-15% of patients who experience treatment failure or relapse^4^ and acute leukemia remains a leading cause of cancer related death in children^2^.

Cellular metabolism plays a role in oncologic processes. In solid tumors, the tumor microenvironment (TME) and nutrient availability influence cancer development, progression, and response to therapy^5–7^. Advancements in mass spectrometry allow identification of metabolites in biological samples, providing key insights into the metabolite composition of tumors. For example, recent studies in renal cell carcinoma (RCC) revealed that nutrient profiles of tumor interstitial fluid closely mirror those of normal kidney tissue, suggesting that RCC cancer cells adapt to the existing metabolic environment within the normal kidney^8^. Thus, metabolic reprogramming of different cancer cell types may be influenced by specific nutrient availabilities within local environments of cancer development.

Emerging evidence in hematologic malignancies also suggests that extrinsic factors like nutrient and metabolite availability may influence leukemic cell metabolism and response to treatment^9,10^. For example, leukemia cells grown in human plasma-like medium (HPLM), which is formulated to match many nutrient levels in blood, exhibit altered nutrient utilization and widespread metabolic rewiring^11^. These environmental changes can affect both drug efficacy and genetic dependencies, as shown by CRISPR screens that revealed context-specific essential genes in HPLM^12^ and by systematic profiling of drug-nutrient interactions^13^. Whether differences in global nutrient availability and utilization influence therapeutic response and outcomes in B-ALL remains an important question.

Here, we utilized quantitative metabolomics to characterize nutrient levels in blood and bone marrow plasma before and after induction chemotherapy in patients with B-ALL with the following categories: hyperdiploidy (HD; favorable genetics), high-risk (HR; unfavorable genetics, patients with induction chemotherapy failure or subsequent relapse), or Philadelphia chromosome-positive (Ph+; unfavorable genetics and received imatinib in addition to chemotherapy during induction). We found baseline subtype-specific alterations in key metabolic pathways, including arginine and creatine metabolism. Further, while most nutrient levels remained stable following chemotherapy, select metabolites showed consistent changes. Comparative analysis with metabolite profiles from renal cell carcinoma (RCC) and non-small cell lung cancer (NSCLC) plasma revealed that pediatric B-ALL has a distinct metabolite signature, notably with enrichment of tricarboxylic acid (TCA) cycle intermediates. Together, these findings reveal distinct metabolic profiles across B-ALL subtypes and identify new avenues for investigation to better understand the underlying biology that may be associated with chemotherapy resistance and poor response to treatment.

## Results

### Sample processing conditions can influence blood plasma metabolite stability

Metabolomics has enabled the study of metabolites in tumors and biological fluids in cancer^14^. These analyses may be sensitive to variability in sample collection and preparation^15,16^. Thus, we first studied the effects of various processing times and collection conditions on metabolite stability in blood using a cohort of patient samples to simulate standard protocols for clinical sample collection and processing used for patients with leukemia. We collected blood samples from healthy donors and either immediately processed and froze the plasma or incubated samples at room temperature or 4°C for various periods of time prior to metabolite extraction for mass spectrometry analysis **(Fig. 1A)**. Using quantitative mass spectrometry, we assessed the stability of polar metabolites detected in plasma under these different processing conditions (**Fig. 1B, Supplementary** Fig. 1**, Supplementary Table 1)**. We found that the majority of detected metabolites remained stable across the tested conditions when compared to immediate processing, with at most 6% of metabolites significantly altered by incubation at room temperature for 1 hour followed by 24 hours at 4°C **(Fig. 1C)**. These data suggest that most measured metabolites remain stable under the tested conditions, with a few notable exceptions. Firstly, hypoxanthine levels increased with extended incubation at 4°C and any period of incubation at room temperature **(Fig. 1D)**, aligning with previous reports that hypoxanthine is secreted by red blood cells during storage at 4°C^16^. Additionally, we noted time-dependent depletion of alpha-ketobutyrate (AKB) with delayed processing **(Fig. 1E)**, which may suggest ongoing cellular uptake of this metabolite during incubation^17,18^. We hypothesized that energy-related metabolites like glucose and lactate would be altered by prolonged processing times due to ongoing metabolism within blood cells *ex vivo*. Interestingly, we found that glucose levels were slightly decreased while lactate levels were increased slightly **(Fig. 1F-G)**. While we cannot fully exclude the possibility of processing variability in patient samples, these data provide a framework to interpret which metabolites may be influenced by handling and help contextualize findings in patient samples. Notably, even processing-sensitive metabolites like hypoxanthine can still yield valuable biological insights when interpreted with awareness of these factors.

**Figure 1.**
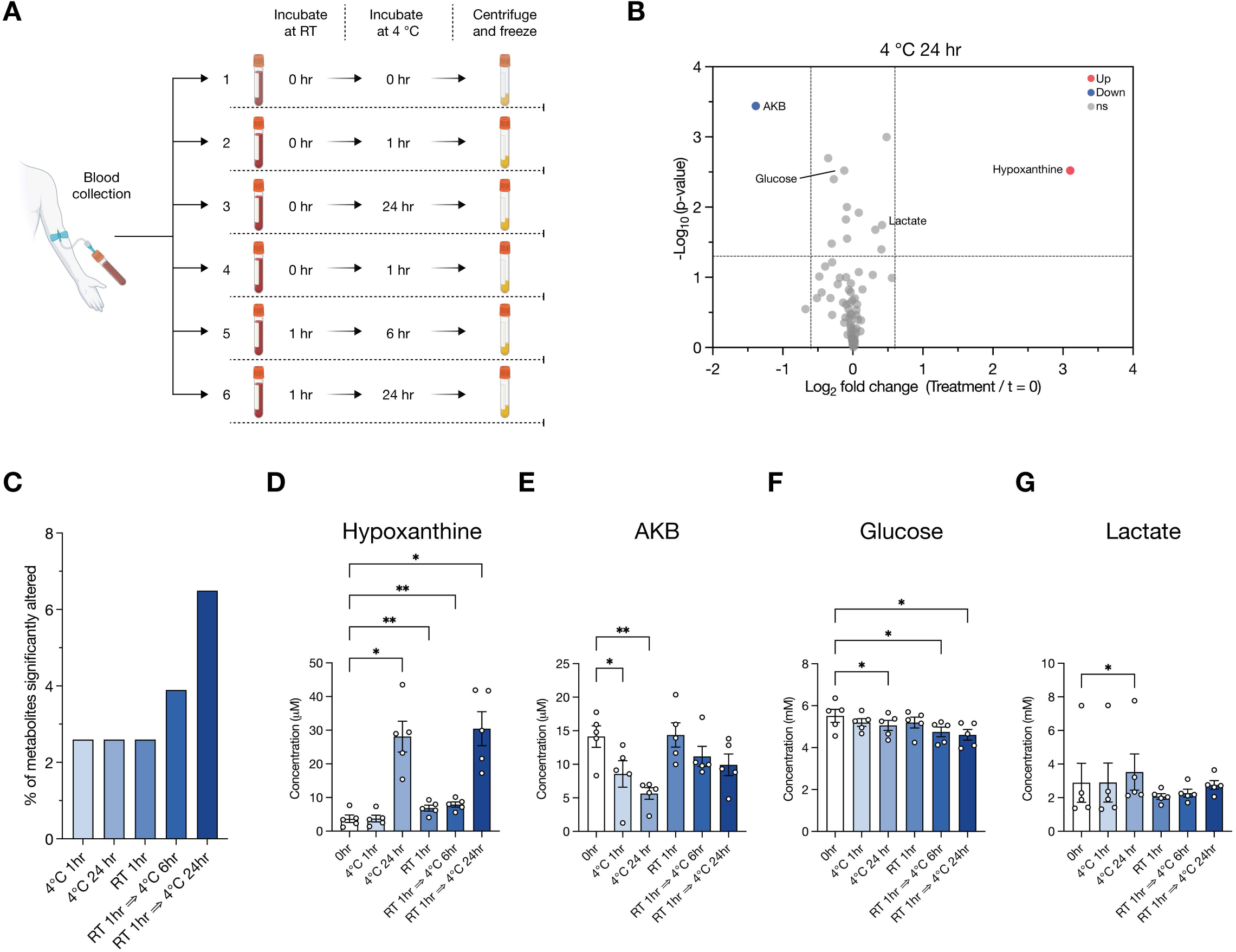
Impact of plasma processing conditions on metabolite stability. **A**, Schematic depicting the experimental design for blood collection and processing conditions. Blood samples were either processed immediately or incubated at room temperature or 4 °C for various durations before plasma extraction. Metabolites in these plasma samples were measured using LC/MS, with quantification performed alongside a dilution series of chemical standards. In total, 77 metabolites were identified and quantified across the different samples. **B**, Volcano plot depicting metabolites that significantly change across processing conditions from n = 6 patients. Cutoffs of | log_2_ fold change | > 0.6 and raw p-value < 0.05 were used to select significantly altered metabolites. AKB: α-ketobutyrate. **C**, Bar plot representing the percent of metabolites in plasma that showed significant changes across different processing conditions compared to immediate processing. Cutoffs of | log_2_ fold change | > 0.6 and raw p-value < 0.05 were used to select significantly altered metabolites. **D-G**, Levels of selected metabolites measured by LC/MS in plasma across processing conditions. Data are presented as mean ± SEM and represent n = 6 patients. P-values were derived from a repeated measures one-way ANOVA with Dunnett’s multiple comparisons test and Geisser-Greenhouse correction (* p < 0.05; **p < 0.01). Statistical comparisons not marked with asterisks were not significant.

### Blood and bone marrow plasma metabolomes are similar in leukemia subtypes

The local nutrient environment can influence cancer cell metabolism^11,19–21^. Thus, we next used quantitative mass spectrometry to investigate how levels of metabolites differ in blood versus bone marrow plasma samples from pediatric patients with B-ALL before and after induction chemotherapy. To this end, we collected blood and bone marrow (BM) plasma samples from patients enrolled on Dana-Farber Cancer Institute clinical trial 05-001 (NCT00400946)^22^. Within this group of patients, we defined three patient subgroups: hyperdiploid (HD; favorable genetics), high-risk (HR; defined as having unfavorable genetics or those who go on to have induction failure, high minimal residual disease after induction, or relapse), and Philadelphia chromosome-positive (Ph+; unfavorable genetics and received imatinib during induction) **(Fig. 2A-B, Supplementary Table 2)**. Blood and BM plasma samples were collected prior to initiation of treatment (day 1), and upon completion of induction chemotherapy (day 32). Blood plasma samples from pediatric healthy donors were used as pre-treatment controls. We performed quantitative metabolomic analyses and compared metabolite concentrations across all samples **(Fig. 2A, Supplementary Table 3)**.

**Figure 2.**
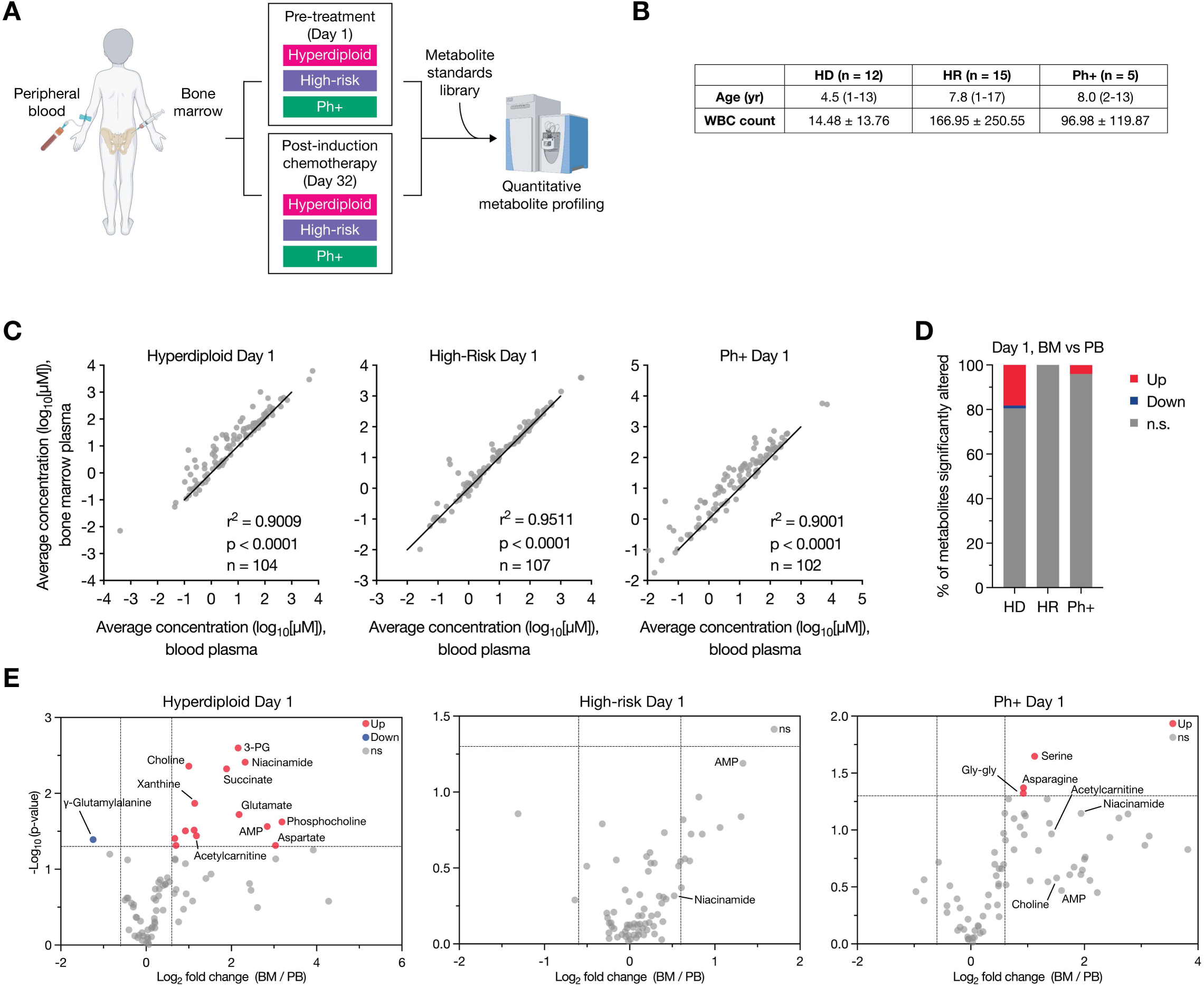
Comparison of bone marrow and blood plasma metabolite composition. **A**, Schematic depicting study design whereby bone marrow or peripheral blood samples were collected from pediatric patients with B-cell acute lymphoblastic leukemia (B-ALL) and metabolites quantified by liquid chromatography/mass spectrometry (LC/MS). Three B-ALL subtypes were analyzed: hyperdiploid (HD), high-risk (HR), and Philadelphia-chromosome translocation positive (Ph+). Samples were collected either on the day of diagnosis/start of induction chemotherapy (day 1) or 32 days post-induction chemotherapy (day 32). **B**, Summary of clinical characteristics of pediatric B-ALL patients by subtype. Table shows the average age at diagnosis and white blood cell (WBC) count on day 1 within each subtype. **C**, Scatter plots of LC/MS measurements of average metabolite concentrations in peripheral blood (PB) or bone marrow (BM) plasma from B-ALL patients on day 1. The number of metabolites measured is indicated in each panel. R and p-values were determined by Pearson correlation. **D**, Bar plot representing the percent of metabolites in plasma that were significantly higher (Up) or lower (Down) in BM compared to PB for each B-ALL subtype on day 1. Cutoffs of | log_2_ fold change | > 0.6 and raw p-value < 0.05 were used to select significantly altered metabolites. n.s.: not significant. **E**, Volcano plots depicting metabolites that significantly change in BM compared to PB for each B-ALL subtype on day 1 (prior to therapy). Cutoffs of | log_2_ fold change | > 0.6 and raw p-value < 0.05 were used to select significantly altered metabolites. For panels C and E, n = 11 (HD PB), n = 12 (HD BM), n = 13 (HR PB), n = 19 (HR BM), or n = 5 (Ph+ PB and Ph+ BM) patient samples.

We first compared metabolite concentrations between blood and BM plasma and found high correlations between the two sampling sites prior to therapy (**Fig. 2C)**. This similarity between blood and bone marrow plasma was also found at day 32 **(Supplementary** Fig. 2A). A subset of analyzed metabolites were present at higher concentrations within the BM plasma compared to blood plasma, particularly in HD and Ph+ subtypes **(Fig. 2D-E, Supplementary** Fig. 2B-C). For instance, niacinamide was enriched in BM in the majority of samples **(Supplementary** Fig. 2D), and other metabolites, including AMP, aspartate, and choline, were also found to be more abundant in BM plasma than in blood plasma in many samples **(Supplementary** Fig. 2E-G). Together, these data highlight that while there are some differences in metabolites between sampling sites, the blood and BM plasma metabolomes are largely comparable to each other before and after induction chemotherapy.

### The blood plasma metabolome of patients with leukemia differs from healthy controls at the time of diagnosis

To investigate whether global metabolomes differ between healthy individuals and patients with leukemia, we compared metabolite levels in blood from healthy donors and B-ALL patients prior to treatment. Principal component analysis (PCA), hierarchical clustering, and statistical analyses all showed a strong separation between metabolite profiles of plasma from patients with leukemia and healthy individuals **(Fig. 3A-C)**. Notably, there was overlap of metabolomes of HR and Ph+ cohorts. Specific metabolites were identified as significantly different between healthy individuals and each leukemia subtype **(Fig. 3D, Supplementary** Fig. 3). Levels of the purine metabolite xanthine were elevated in all leukemia samples compared to healthy controls, regardless of subtype **(Fig. 3E)**, which may reflect increased turnover of leukemic cells and subsequent xanthine production^23^. Uric acid levels were significantly increased in HR samples compared to all other subtypes and healthy donors, which may also be a marker of increased cell turnover **(Fig. 3F**. Interestingly, arginine concentrations were lower in HD samples compared to other subtypes and healthy controls **(Fig. 3G)**, whereas levels of asymmetric dimethylarginine (ADMA) were significantly higher **(Fig. 3H)**, suggesting that these specific leukemia categories differ in arginine metabolism. Aspartate, a key substrate in the synthesis of purines and pyrimidines, was elevated in all leukemic samples compared to healthy donors **(Fig. 3I)**. Finally, we observed that guanidinoacetate (GAA) concentrations were lower in plasma from all patients with leukemia compared to healthy controls, with the most pronounced decrease noted in HR and Ph+ subtypes **(Fig. 3J)**. Though GAA is the direct precursor of creatine, we did not observe significant differences in creatine levels across healthy donors and patients with leukemia **(Fig. 3K)**. Taken together, these data reveal that the blood plasma metabolome of leukemia patients is different from healthy counterparts at the time of diagnosis and that subtype-specific differences exist.

**Figure 3.**
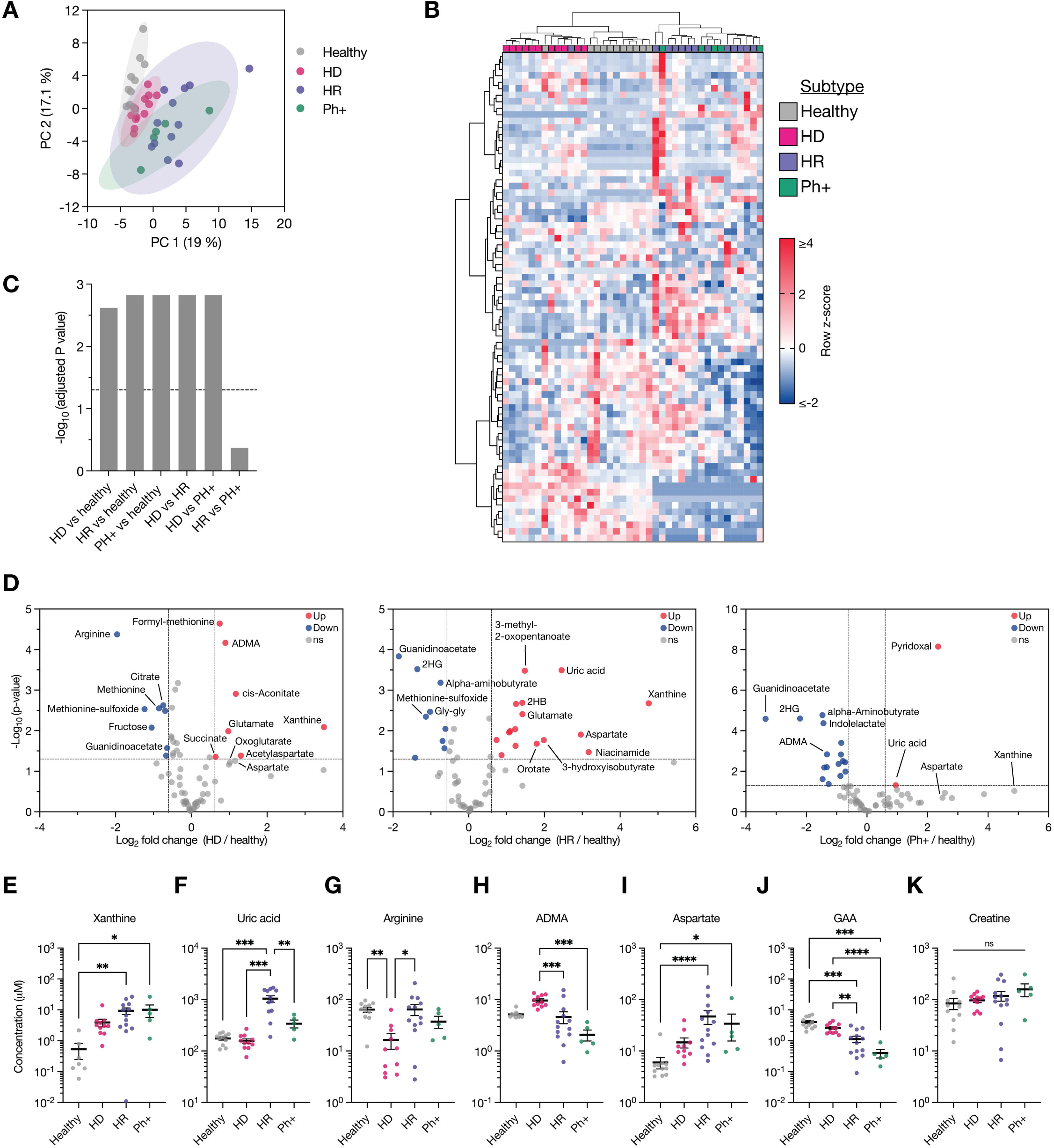
**Comparison of plasma nutrient profiles across B-ALL subtypes and healthy controls. A-B**, Principal component analysis (PCA) (A) or hierarchical clustering (B) of metabolites measured in peripheral blood plasma from healthy donors or B-ALL patients. Three B-ALL subtypes were analyzed: hyperdiploid (HD), high-risk (HR), and Philadelphia-chromosome translocation positive (Ph+). Data represent n = 11 (healthy), n = 11 (HD), n = 13 (HR), or n = 5 (Ph+) patient samples. Data presented within each row of the heatmap were z-score normalized. **C**, Bar plot showing false discovery rate (FDR)-corrected p-values from pairwise PERMANOVA analysis of the PCA plot in (A). The dotted line indicates a significance threshold of p = 0.05. **D**, Volcano plots depicting metabolites that significantly change in concentration between B-ALL patients on day of diagnosis and healthy patients. Cutoffs of | log_2_ fold change | > 0.6 and raw p-value < 0.05 were used to select significantly altered metabolites. **E-K**, Levels of selected metabolites measured by LC/MS in peripheral blood plasma from healthy patients or B-ALL patients on day of diagnosis. Data are presented as mean ± SEM. p-values were derived from a Kruskal-Wallis test with Dunn’s multiple comparisons correction (xanthine, aspartate, arginine, ADMA, creatine) or from a Brown-Forsythe and Welch ANOVA with Dunnett’s T3 multiple comparisons test (uric acid, GAA) (* p < 0.05; **p < 0.01; ***p < 0.001; ****p < 0.0001; ns, not significant). For panels D-K, n = 11 (healthy), n = 11 (HD), n = 13 (HR) or n = 5 (Ph+) patient samples. Statistical comparisons not marked with asterisks were not significant. 2HG: 2-hydroglutarate; ADMA: asymmetric dimethylarginine; GAA: guanidinoacetic acid gly-gly: glycyl-glycine.

Type 1 protein arginine methyltransferases (PRMTs) are enzymes that methylate arginine residues to produce ADMA^24–26^. To explore whether increased ADMA levels in HD samples might reflect enhanced activity of PRMTs within B-ALL, we queried the Cancer Dependency Map Project (DepMap) for PRMT gene expression in pediatric B-ALL cell lines compared to other malignancies. We found that pediatric B-ALL cell lines are particularly dependent on the type I PRMTs *CARM1* (also known as *PRMT4*), *PRMT1*, and *PRMT8*, as well as the type II *PRMT9*. This correlates with an elevated expression of a subset of PRMTs in B-ALL compared to other cancers **(Supplementary** Fig. 4A-B). These findings suggest that protein arginine methylation may represent a leukemia-selective metabolic vulnerability and might warrant further investigation as a potential therapeutic target in B-ALL.

Previous studies have shown that certain leukemias, such as EVI1-positive acute myeloid leukemia (AML), upregulate mitochondrial creatine kinase (CKMT1) and rely on creatine metabolism for energy homeostasis^27^. Given lower levels of the creatine precursor GAA in leukemia subtypes compared to healthy controls, we again queried DepMap for expression of genes that are part of the creatine synthesis pathway. Pediatric B-ALL cell lines upregulate *GAMT* expression compared to other cell lines **(Supplementary** Fig. 4C), although they do not display increased dependency on any creatine synthesis pathway genes. However, the limited number of pediatric B-ALL models currently available in DepMap may restrict assessment of dependency on creatine metabolism in this disease.

### Induction chemotherapy influences a discrete subset of metabolic pathways in leukemia samples

Next, we sought to understand whether metabolite concentrations are influenced by induction chemotherapy. We performed comparative analyses of metabolites found in blood or BM plasma from leukemia patients prior to and after completion of induction chemotherapy. Within leukemia subtypes, metabolite concentrations were highly correlated between samples collected before and after treatment **(Fig. 4A, Supplementary** Fig. 5A), with most metabolites showing no significant changes at day 32 compared to day 1 **(Fig. 4B, Supplementary** Fig. 5B). However, several noteworthy differences emerged in our analysis. Specifically, asparagine levels were significantly lower in both blood and BM plasma of HD patients following induction chemotherapy across all subtypes, reflecting the use of asparaginase during induction chemotherapy **(Fig. 4C-D, Supplementary** Fig. 5C-D). In many post-treatment samples, asparagine levels fell below the limit of detection by LC/MS, which limited statistical power to detect significant differences despite a consistent trend of depletion (**Fig.4D**). As asparagine is hydrolyzed to aspartate, we next evaluated aspartate levels and found mild non-significant increases in HD plasma samples and unchanged levels in HR and Ph+ samples **(Supplementary** Fig. 5E-F). We also observed a significant reduction in the levels of the dipeptide glycyl-glycine (Gly-Gly) after chemotherapy in both blood and BM plasma **(Fig. 4C and 4E, Supplementary** Fig. 5G), with post-treatment levels falling below the detection limit in several cases. Xanthine levels declined in all B-ALL subtypes after chemotherapy **(Fig. 4G, Supplementary** Fig. 5H), which may reflect decreasing cell turnover as leukemic burden is reduced. Furthermore, ADMA decreased across all leukemia subtypes, with a more pronounced reduction observed in HD and HR patients **(Fig. 4G, Supplementary** Fig. 5I). Collectively, these findings suggest that while chemotherapy does not extensively alter the overall nutrient environment, it does impact a discrete subset of metabolites. Deeper understanding of these altered pathways in leukemia subtypes may provide insights into mechanisms underlying treatment failure and relapse.

**Figure 4.**
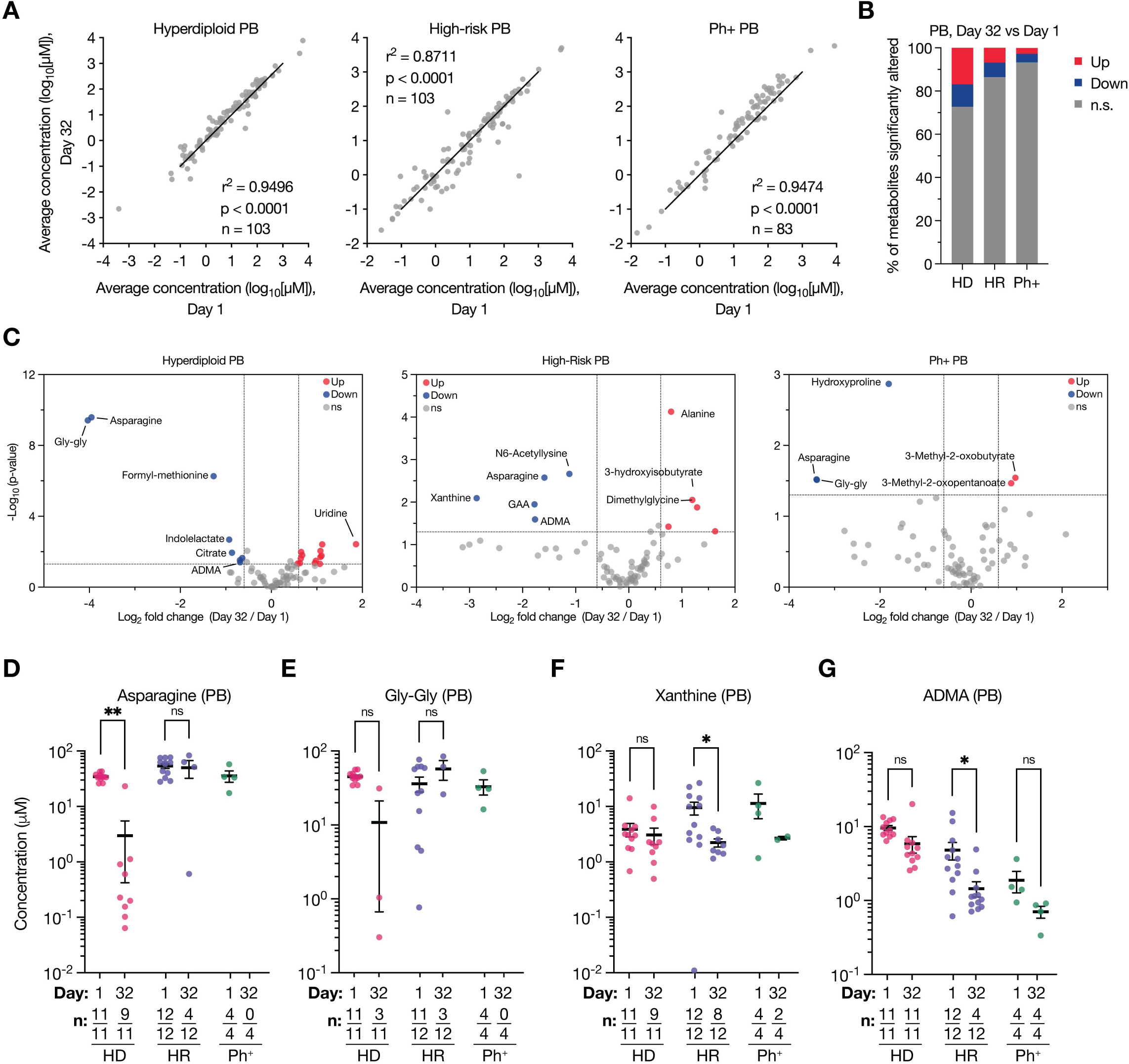
Analysis of metabolite changes in peripheral blood plasma of B-ALL patients pre-and post-induction chemotherapy. **A**, Scatter plots of metabolite concentrations measured by LC/MS in peripheral blood (PB) plasma from B-ALL patients on day 32 post-induction chemotherapy (day 32) compared to the day of diagnosis (day 1). The number of metabolites measured is indicated in each panel. r and p-values were determined by Pearson correlation. **B**, Bar plot representing the percent of metabolites in peripheral blood plasma that were significantly higher (Up) or lower (Down) on day 32 compared to day 1. Cutoffs of | log2 fold change | > 0.6 and raw p-value < 0.05 were used to select significantly altered metabolites. n.s.: not significant. **C**, Volcano plots depicting metabolites that significantly change in concentration in in PB plasma between day 1 and day 32 post-induction chemotherapy. Cutoffs of | log_2_ fold change | > 0.6 and raw p-value < 0.05 were used to select significantly altered metabolites. **D-H**, Levels of selected metabolites measured by LC/MS in PB plasma from paired B-ALL patients on day 1 or day 32 post-induction chemotherapy. Data are presented as mean ± SEM and p-values were derived from a Wilcoxon matched-pairs signed rank test with Holm-Sidak’s multiple comparisons correction (* p < 0.05; **p < 0.01; ns, not significant). n values indicate the number of samples with detectable metabolite levels (numerator) out of the total number of samples analyzed (denominator). ADMA: asymmetric dimethylarginine; GAA: guanidinoacetic acid; Gly-Gly: glycyl-glycine.

### The leukemia plasma metabolome is markedly different from RCC and NSCLC interstitial fluid metabolomes

Finally, we compared plasma metabolite concentrations in samples from pediatric patients with B-ALL with those from adult patients with renal cell carcinoma (RCC) or non-small cell lung cancer (NSCLC). Specifically, we performed comparative analysis of blood plasma metabolites from healthy donors and B-ALL patients prior to treatment versus previously published plasma metabolite concentrations from RCC and NSCLC patients^21,8^. PCA and hierarchical clustering indicated some similarities among groups **(Fig. 5A, Supplementary** Fig. 6); however, statistical analyses revealed a high degree of separation in overall metabolite composition across groups **(Fig. 5B)**. Notably, NSCLC and RCC samples were more similar to each other than to leukemia samples. Several metabolites differed significantly between healthy, B-ALL, NSCLC, and RCC samples. For instance, hypoxanthine and ADMA were both elevated in B-ALL compared to NSCLC and RCC samples **(Fig. 5D,E)**. Many intermediates of the tricarboxylic acid (TCA) cycle and related metabolites, including pyruvate, oxoglutarate, fumarate, succinate, glutamate, and aspartate, were elevated in B-ALL relative to NSCLC and RCC, with the exception of citrate **(Fig. 5F-L)**. In line with this observation, analysis of the Depmap further highlighted the TCA cycle as a dependency in pediatric B-ALL cell lines compared to other tumor types **(Supplementary** Fig. 7). Of note, cystine levels were lower in B-ALL compared to NSCLC and RCC, and were closer to that observed in healthy controls **(Fig. 5M)**. Overall, these findings demonstrate that patients with leukemia and those with solid tumors have largely different metabolomes and highlights the need for further studies to identify the role of altered metabolic pathways in the development, progression, and response to treatment of hematologic malignancies.

**Figure 5.**
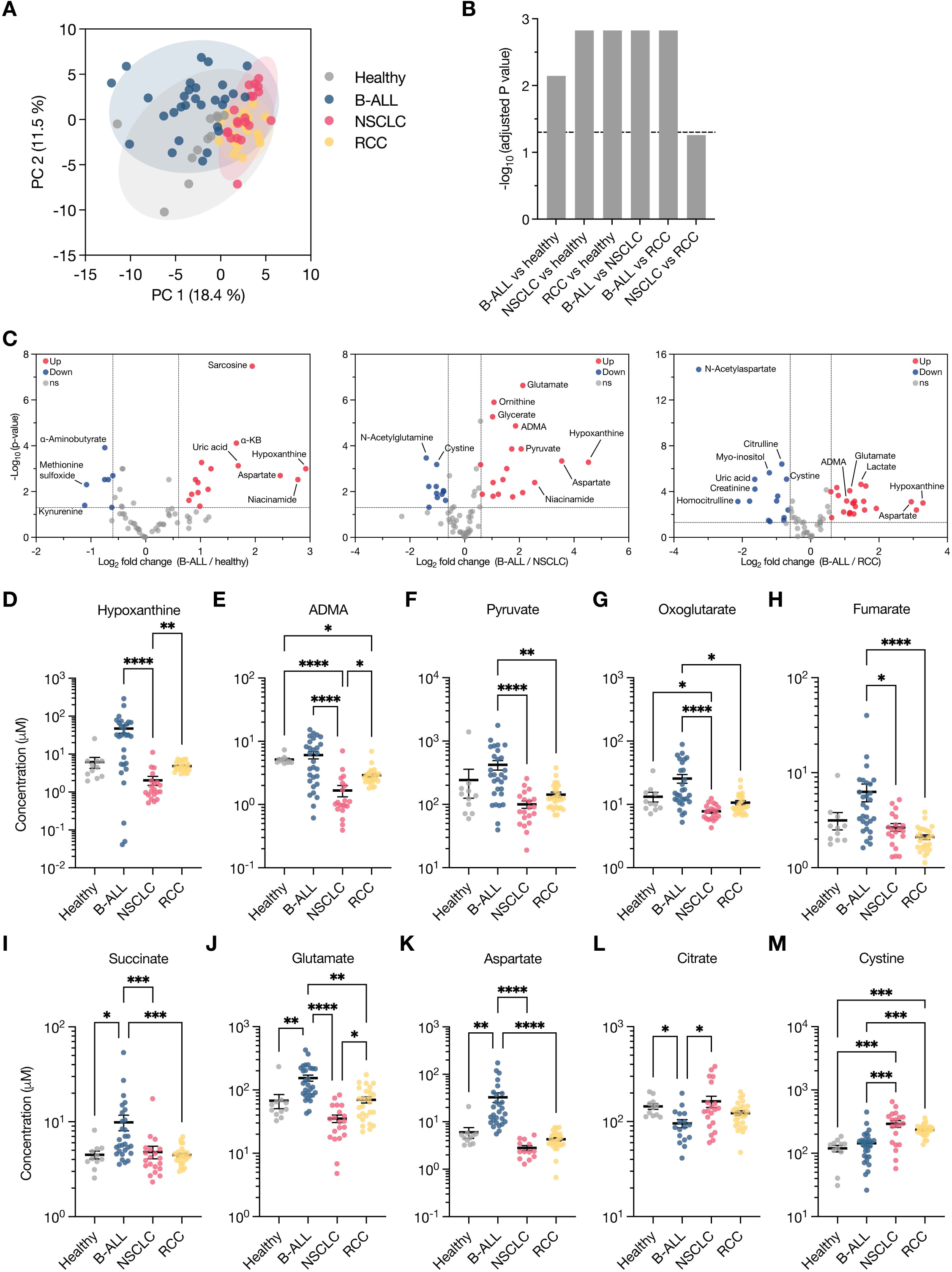
Comparison of peripheral blood plasma metabolite composition in B-ALL, NSCLC, and RCC patients. **A**, Principal component analysis (PCA) of metabolites measured in peripheral blood plasma from healthy donors and patients with B-ALL (day of diagnosis), non-small cell lung cancer (NSCLC), or renal cell carcinoma (RCC). Data represent n = 11 (healthy), n = 29 (B-ALL), n = 20 (NSCLC), and n = 27 (RCC) patient samples. For this analysis, B-ALL subtypes were combined. **B**, Bar plot showing false discovery rate (FDR)-corrected p-values from pairwise PERMANOVA analysis of the PCA plot in (A). The dotted line indicates a significance threshold of p = 0.05. **C**, Volcano plots depicting metabolites that significantly change in concentration between B-ALL patients (day of diagnosis) and healthy donors, NSCLC patients, or RCC patients. Cutoffs of | log_2_ fold change | > 0.6 and raw p-value < 0.05 were used to select significantly altered metabolites. Data represent n = 11 (healthy), n = 29 (B-ALL), n = 20 (NSCLC), and n = 27 (RCC) patient samples. **D-M**, Levels of selected metabolites measured by LC/MS in peripheral blood plasma from healthy donors and patients bearing B-ALL (day of diagnosis), NSCLC, or RCC. Data are presented as mean ± SEM and p-values were derived from a Kruskal-Wallis test with Dunn’s multiple comparisons correction (* p < 0.05; **p < 0.01; ***p < 0.001; ****p < 0.0001; ns, not significant). Statistical comparisons not marked with asterisks were not significant. For all panels, n = 11 (healthy), n = 29 (B-ALL), n = 20 (NSCLC), or n = 27 (RCC) patients. α-KB: alpha-ketobutyrate; ADMA: asymmetric dimethylarginine.

## Discussion

This study reveals that pediatric B-ALL is associated with a distinct plasma metabolome characterized by subtype-specific alterations. While overall nutrient levels remain relatively stable across compartments and after induction chemotherapy, several metabolites exhibit consistent and potentially clinically relevant changes. For instance, specific alterations in arginine and ADMA levels in hyperdiploid B-ALL, and the pronounced depletion of asparagine and Gly-Gly following treatment, point to metabolic shifts that may reflect underlying disease biology or treatment response. These findings highlight discrete and potentially biologically meaningful metabolic features of B-ALL that could inform future studies aimed at biomarker discovery and therapeutic intervention.

Processing conditions and timing of clinical samples collection vary across research groups and studies. Our data demonstrate that plasma processing techniques influence the stability of some metabolites. Specifically, longer incubation periods prior to processing and freezing were correlated with higher levels of hypoxanthine and lactate and lower glucose levels. These results are consistent with prior studies of red blood cell metabolism demonstrating ongoing hypoxanthine secretion and glucose metabolism in samples stored at 4°C^15,16^. These data detailing the stability of various metabolites under different processing conditions will serve as a resource for other investigators who are performing metabolomic studies using clinical samples.

Interestingly, we found that nutrient profiles detected by mass spectrometry are overall similar between blood and BM plasma samples, both before and after induction chemotherapy. Similarly, Schraw *et al* previously found considerable overlap in measured metabolites across serum, bone marrow, and CSF samples of pediatric B-ALL patients^10^. These data suggest that peripheral blood could be used in place of more invasive bone marrow sampling for metabolite biomarker studies in pediatric B-ALL. However, it is important to note that other groups have reported more pronounced differences in metabolites detected in the BM versus blood plasma of patients with leukemia^28–30^. These discrepancies may arise from several factors, including differences in induction chemotherapy regimens, sample processing protocols, and technical variations in metabolite measurement across studies. A more extensive, standardized approach — encompassing uniform chemotherapy protocols and consistent processing methods — will better clarify whether these differences are methodological or biologically driven. Comparisons involving healthy BM and blood plasma will also provide an essential baseline for interpreting leukemia-specific metabolic shifts.

Asparaginase is an enzyme that degrades the amino acid asparagine and is a cornerstone of ALL treatment^31^. Children with ALL who have suboptimal exposure to asparaginase, often due to toxicity or silent inactivation, are at an increased risk of relapse^32–34^. In line with its mechanism, we found that asparagine levels were markedly reduced across all samples following induction chemotherapy. This indicates that asparaginase effectively depleted its target metabolite even in patients with poor-risk subtypes. Interestingly, Gly-Gly, a dipeptide not directly targeted by therapy, showed a similarly consistent pattern of depletion. While the biological relevance of this finding remains unclear, the parallel reduction in Gly-Gly suggests additional, underexplored shifts in peptide metabolism during treatment and highlights the value of untargeted metabolomics in uncovering unexpected therapy-associated metabolite changes.

Perhaps unsurprisingly, our analyses found that there are significant differences in metabolite profiles between healthy and leukemia patients. Notably, the metabolomes of HR and Ph+ leukemia largely clustered together, consistent with their shared status as clinically high-risk subtypes. In contrast, HD samples exhibited distinct features, including significantly lower arginine levels and higher concentrations of ADMA. These findings suggest a shift in arginine metabolism in HD B-ALL, potentially driven by increased PRMT activity, as pediatric B-ALL cell lines show selective dependency on the type I PRMT genes *CARM1*, *PRMT1*, and *PRMT8*, as well as the type II *PRMT9*. Our findings align with a recent study that also identified altered arginine metabolism in pediatric B-ALL^28^; however, while their work focused on suppression of argininosuccinate synthase (ASS1) and disruption of the urea cycle, our data point to a distinct mechanism involving ADMA accumulation and PRMT activity, particularly in HD. Together, these findings suggest that arginine metabolism is disrupted through multiple, potentially subtype-specific pathways in B-ALL, and further investigation is warranted to understand how these metabolic alterations influence disease progression and therapeutic response.

When comparing the B-ALL blood metabolome to that of RCC and NSCLC, we observe a distinct enrichment of TCA cycle intermediates, consistent with DepMap analyses revealing a heightened TCA cycle dependence in B-ALL cells. Although factors such as patient age, fasting status, and processing protocols may complicate cross-cancer comparisons, these results highlight distinctions between hematologic and solid tumors. Interestingly, prior studies suggest that solid tumors like RCC do not strongly modulate their local nutrient environment^8^. In contrast, the observed differences between healthy and leukemia plasma in our study raise the possibility that leukemic cells have a larger impact on circulating nutrient levels, possibly due to their large numbers in blood. This distinction emphasizes the importance of tumor type in shaping the nutrient microenvironment and has implications for understanding cancer metabolism and host interactions.

Taken together, this work informs nutrient levels in pediatric B-ALL and highlights how nutrient availability can differ between hematologic malignancies and solid tumors. By identifying distinct metabolic alterations across B-ALL subtypes, we provide a foundation for exploring metabolic vulnerabilities specific to pediatric leukemias that might be shaped by the nutrient environment. Future studies are needed to validate the relevance of these metabolites as biomarkers or therapeutic targets and to investigate their broader significance in predicting treatment responses.

## Supporting information

Supplementary Table 1 - Metabolite stability experiment

Supplementary Table 2 - Patient characteristics

Supplementary Table 3 - Metabolite concentrations

Supplementary Table 4 - DepMap pediatric B-ALL cell lines

## Acknowledgements

We thank members of the Vander Heiden and Pikman laboratories for helpful discussions. K.L.A. was supported by the National Science Foundation (DGE-1122374) and National Institutes of Health (NIH) (F31CA271787, T32GM007287). A.A. received support as a Howard Hughes Medical Institute (HHMI) Medical Research Fellow. B.K. acknowledges support from the NIH (R01CA249185). M.G.V.H. acknowledges support from the MIT Center for Precision Cancer Medicine, the Ludwig Center at MIT, Stand Up to Cancer, the Lustgarten foundation, an HHMI Faculty Scholars grant, and the NIH (R35CA242379, P30CA1405141). Y.P. received support from the NIH (K08CA222684) and Hyundai Hope on Wheels. This work was supported by the Pediatric Cancer Dependencies Accelerator of the Broad Institute, Dana-Farber Cancer Institute, and St. Jude Children’s Research Hospital: peddep.org.

## Authorship Contributions

Conceptualization: K.L.A., A.A., M.G.V.H., Y.P.; Methodology: K.L.A., A.A., M.G.V.H., Y.P.; Investigation: K.L.A., A.A., M.B.M., D.B., T.K., M.W., C.R.M.W., B.D.; Resources: M.H.H., B.K., L.B.S., M.G.V.H., Y.P.; Writing – Original Draft: K.L.A., D.B., M.G.V.H., Y.P.; Writing – Review & Editing: All authors; Supervision: M.G.V.H., Y.P.; Funding Acquisition: M.G.V.H., Y.P.

## Methods

### Patient characteristics and sample collection

Paired bone marrow and peripheral blood plasma specimens from 32 children diagnosed with B-ALL, enrolled on clinical trial DFCI 05-001 (NCT00400946) were collected at Boston Children’s Hospital/Dana-Farber Cancer Institute (Boston, MA)^35^. Written informed consent was obtained in accordance with Institutional Review Board guidelines. Patients received induction chemotherapy according to the schema outlined in Appendix Table 2 from Vrooman et al^35^. Patient characteristics are given in Supplementary Table 2. Bone marrow and peripheral blood specimens were obtained on day 1 of treatment and at the end of induction therapy (day 32). Blood and bone marrow samples were collected into EDTA-coated tubes, and plasma was separated by centrifugation at 800 x g for 10 min at 4°C. Samples were snap-frozen and stored at −80 °C prior to analysis by liquid chromatography/mass spectrometry (LC/MS). Plasma metabolite values from renal cell carcinoma (RCC) or non-small cell lung cancer (NSCLC) are previously reported^8^.

For studying processing conditions in healthy patients, peripheral blood samples were collected from consented healthy pediatric and adolescent/young adult patients who were being evaluated in a cancer predisposition clinic and did not have cancer. After collection into EDTA-coated tubes, samples were either processed immediately or subjected to delayed processing as described in Figure 1A. Plasma was isolated by centrifugation at 800 x g for 10 minutes at 4 °C and then snap-frozen and stored at −80 °C until analysis by LC/MS.

## Mass spectrometry metabolite measurements

### Quantification of metabolite levels in biological fluids

Metabolite quantification in human fluid samples was performed as described previously^36^. In brief, 5 μL of sample or external chemical standard pool (ranging from ∼5 mM to ∼1 μM) was mixed with 45 μL of acetonitrile:methanol:formic acid (75:25:0.1) extraction mix including isotopically labeled internal standards. All solvents used in the extraction mix were HPLC grade. Samples were vortexed for 15 min at 4 °C and insoluble material was sedimented by centrifugation at 16,000 x g for 10 min at 4 °C. 20 μL of the soluble polar metabolite extract was taken for LC/MS analysis (described below). Following analysis by LC/MS, metabolite identification was performed with XCalibur 2.2 software (Thermo Fisher Scientific) using a 5 ppm mass accuracy and a 0.5 min retention time window. For metabolite identification, external standard pools were used for assignment of metabolites to peaks at given m/z and retention time. Absolute metabolite concentrations were determined as published^36^.

### LC/MS analysis

Metabolite profiling was conducted on a QExactive benchtop orbitrap mass spectrometer equipped with an Ion Max source and a HESI II probe, which was coupled to a Dionex UltiMate 3000 HPLC system (Thermo Fisher Scientific). External mass calibration was performed using the standard calibration mixture every 7 days. An additional custom mass calibration was performed weekly alongside standard mass calibrations to calibrate the lower end of the spectrum (m/z 70-1050 positive mode and m/z 60-900 negative mode) using the standard calibration mixtures spiked with glycine (positive mode) and aspartate (negative mode). 2 μL of each sample was injected onto a SeQuant® ZIC®-pHILIC 150 x 2.1 mm analytical column equipped with a 2.1 x 20 mm guard column (both 5 mm particle size; EMD Millipore). Buffer A was 20 mM ammonium carbonate, 0.1% ammonium hydroxide; Buffer B was acetonitrile. The column oven and autosampler tray were held at 25 °C and 4 °C, respectively. The chromatographic gradient was run at a flow rate of 0.150 mL min-1 as follows: 0-20 min: linear gradient from 80-20% B; 20-20.5 min: linear gradient form 20-80% B; 20.5-28 min: hold at 80% B. The mass spectrometer was operated in full-scan, polarity-switching mode, with the spray voltage set to 3.0 kV, the heated capillary held at 275 °C, and the HESI probe held at 350 °C. The sheath gas flow was set to 40 units, the auxiliary gas flow was set to 15 units, and the sweep gas flow was set to 1 unit. MS data acquisition was performed in a range of m/z = 70–1000, with the resolution set at 70,000, the AGC target at 1×10^6^, and the maximum injection time at 20 msec.

### DepMap dependency and expression analyses

Gene dependency and expression data were obtained from the DepMap 24Q4 release. For gene-level analyses, violin plots were generated to compare gene dependencies and gene expression values (log_2_ TPM + 1) between pediatric B-cell leukemia cell lines and other cancer cell lines. For dependency data, comparisons were made between 10 pediatric B-cell leukemia cell lines and 1,168 other cancer cell lines. For expression data, comparisons included 17 pediatric B-cell leukemia cell lines and 1,656 other cancer cell lines. Statistical significance was assessed using two-tailed Mann-Whitney tests. For pathway-level analysis, KEGG pathway enrichment was performed using gene set enrichment analysis, comparing pediatric B-lymphoblastic leukemia/lymphoma cell lines (n = 13) to all other cancer types (n = 1,137), NSCLC (n = 98), or RCC (n = 29). Normalized enrichment scores were visualized using volcano plots, with the TCA cycle pathway highlighted. Pediatric B-cell leukemia cell lines used in these analyses and their characteristics are listed in Supplementary Table 4.

### Quantification and statistical analysis

Sample sizes, reproducibility and statistical tests used for each figure are denoted in the figure legends. All graphs were generated using GraphPad Prism 10 (GraphPad Software).

After determining the concentration of each metabolite in each plasma sample, all multivariate statistical analysis on the data was performed using Metaboanalyst 6.0^37^. Metabolite concentrations were autoscaled and metabolites that contained greater than 50% missing values were removed prior to analysis. For the remainder of the metabolites, missing values were replaced using 1/5^th^ of the lowest positive value. All further statistical information is described in the figure legends.

## Data Availability

All metabolite concentration data generated and analyzed during this study are included in this published article and its supplementary information files. Further data required to re-analyze the findings are available from the corresponding author upon reasonable request.

## Illustrations

Experimental schema and illustrative models were generated using BioRender (https://biorender.com/).

---

**Supplementary Figure 1.**
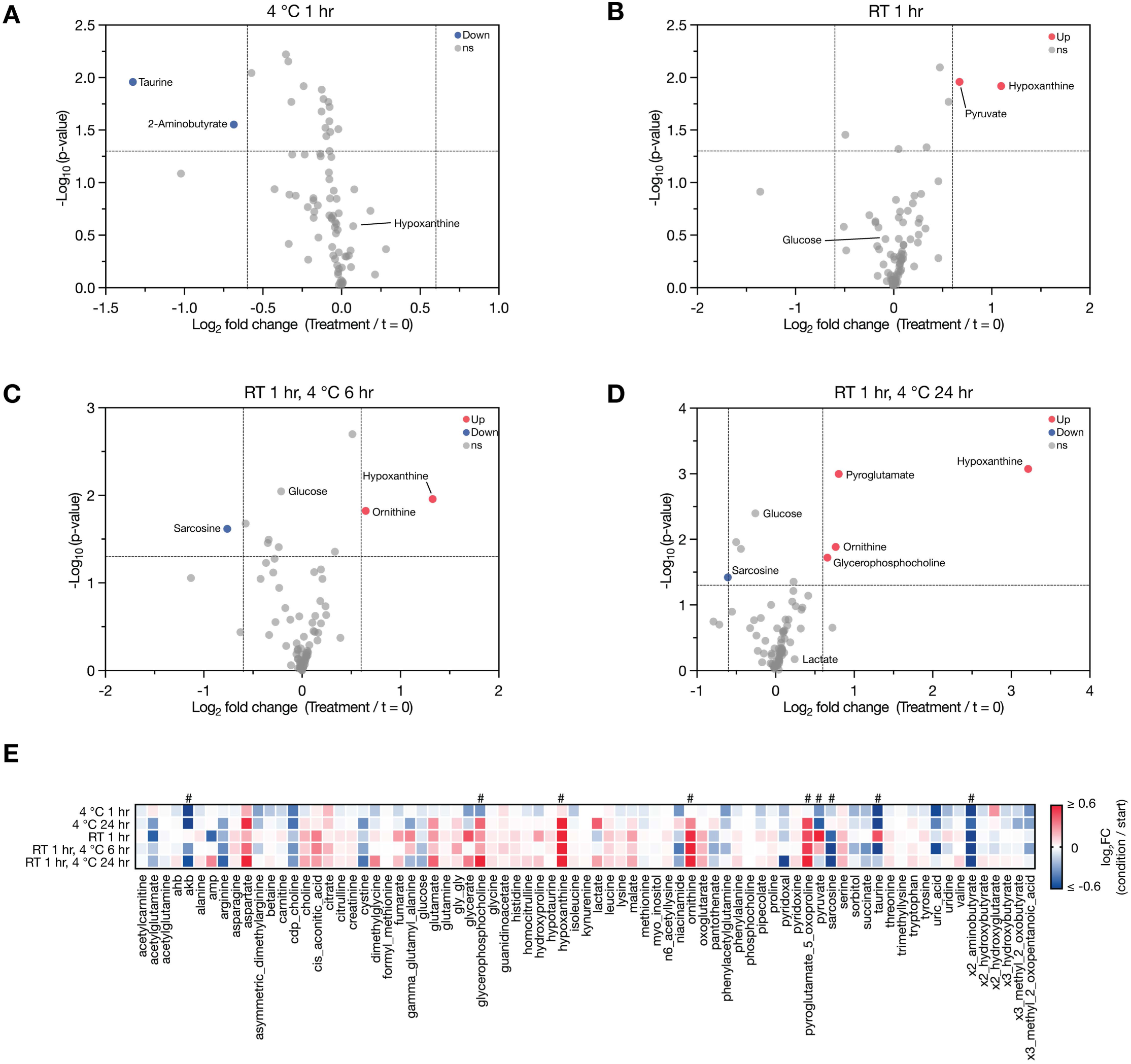
**Analysis of metabolite changes due to plasma processing conditions. A-D**, Volcano plots depicting metabolites that significantly change during different processing conditions from n = 6 patients as indicated. Cutoffs of | log_2_ fold change | > 0.6 and raw p-value < 0.05 were used to select significantly altered metabolites. **E**, Heatmap depicting the average log_2_ fold change in metabolite concentrations measured in plasma across all processing conditions compared to the initial concentration. Scale bar indicates the ranges of values shown. Metabolites that showed significant changes (| log_2_ fold change | > 0.6 and raw p-value < 0.05) in any condition are marked with a # symbol.

**Supplementary Figure 2.**
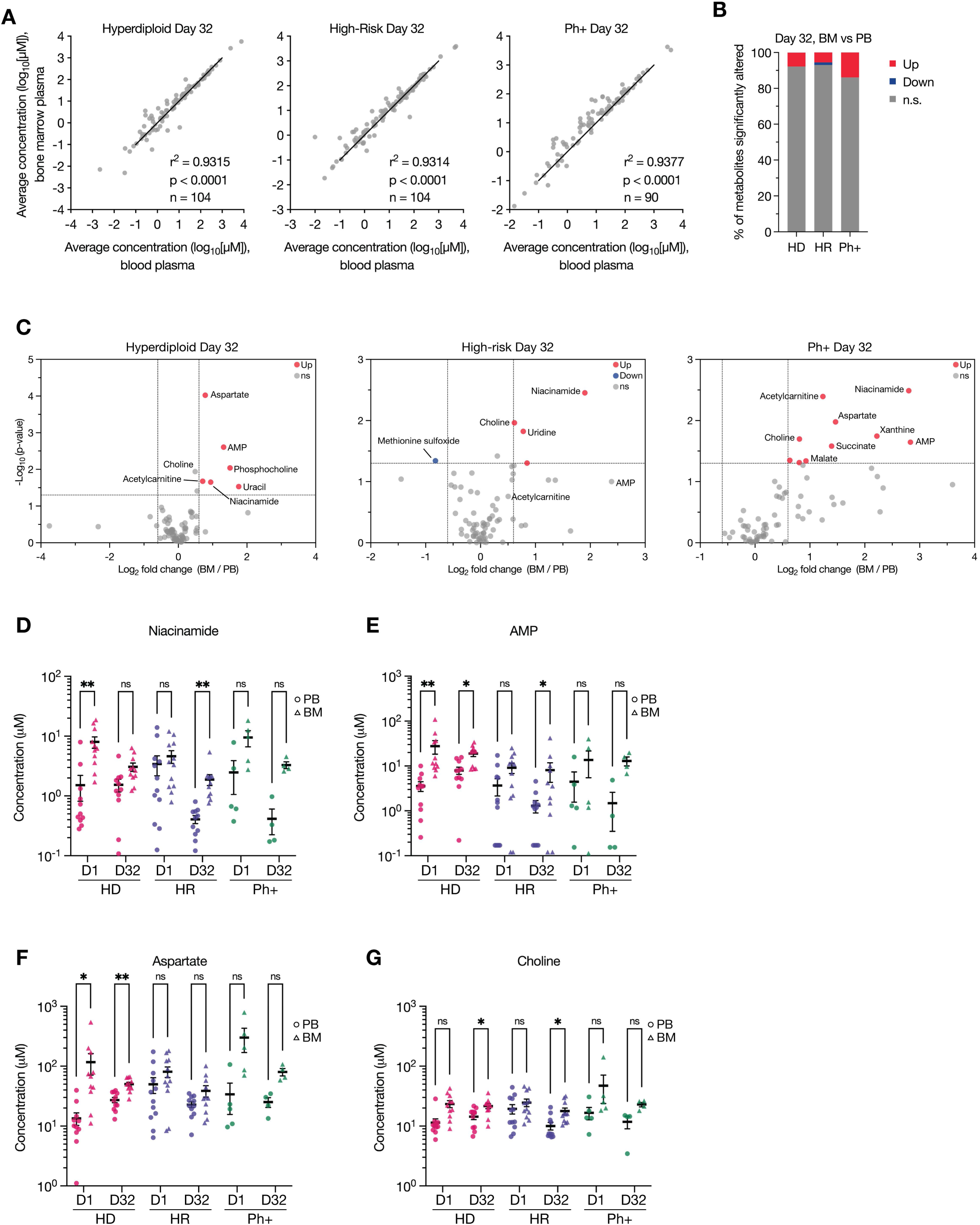
Comparison of bone marrow and blood plasma metabolites post-induction chemotherapy. **A**, Scatter plots of LC/MS measurements of metabolite concentrations in peripheral blood (PB) or bone marrow (BM) plasma from B-ALL patients on day 32 post-induction chemotherapy. The number of metabolites measured is indicated in each panel. r and p-values were determined by Pearson correlation. **B**, Bar plot representing the percent of metabolites in plasma that were significantly higher (Up) or lower (Down) in BM compared to PB plasma for each B-ALL subtype on day 32 post-chemotherapy induction. Cutoffs of | log2 fold change | > 0.6 and raw p-value < 0.05 were used to select significantly altered metabolites. n.s.: not significant. **C**, Volcano plots depicting metabolites that significantly change in BM compared to PB plasma for each B-ALL subtype on day 32 post-chemotherapy induction. Cutoffs of | log_2_ fold change | > 0.6 and raw p-value < 0.05 were used to select significantly altered metabolites. **D-G**, Levels of selected metabolites measured by LC/MS in peripheral blood (PB, circles) or bone marrow (BM, triangles) plasma from B-ALL patients on day 1 (D1) or day 32 (D32) post-induction chemotherapy. Data are presented as mean ± SEM and p-values were derived from a Wilcoxon matched-pairs signed rank test with Holm-Sidak’s multiple comparisons correction (* p < 0.05; **p < 0.01). For panels A and C, n = 12 (HD PB), n = 11 (HD BM), n = 14 (HR PB), n = 17 (HR BM), or n = 4 (Ph+ PB), or n = 5 (Ph+ BM) patient samples. For panels D-H, n = 11 (HD D1 and HD D32), n = 12 (HR D1), n = 11 (HR D32), n = 5 (Ph+ D1), or n = 4 (Ph+ D32) paired patient samples.

**Supplementary Figure 3.**
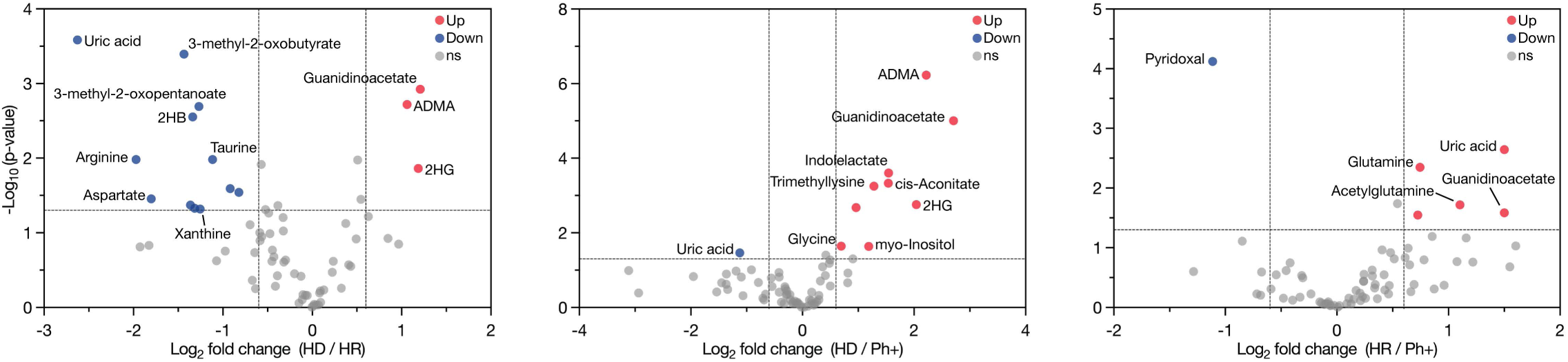
Comparison of plasma nutrient levels across B-ALL subtypes. Volcano plots depicting metabolites that significantly change in plasma concentration between different B-ALL subtypes. Cutoffs of | log_2_ fold change | > 0.6 and raw p-value < 0.05 were used to select significantly altered metabolites. Data represent n = 11 (HD), n = 13 (HR) or n = 5 (Ph+) patient samples. 2HB: 2-hydroxybutyrate; 2HG: 2-hydroxyglutarate; ADMA: asymmetric dimethylarginine.

**Supplementary Figure 4.**
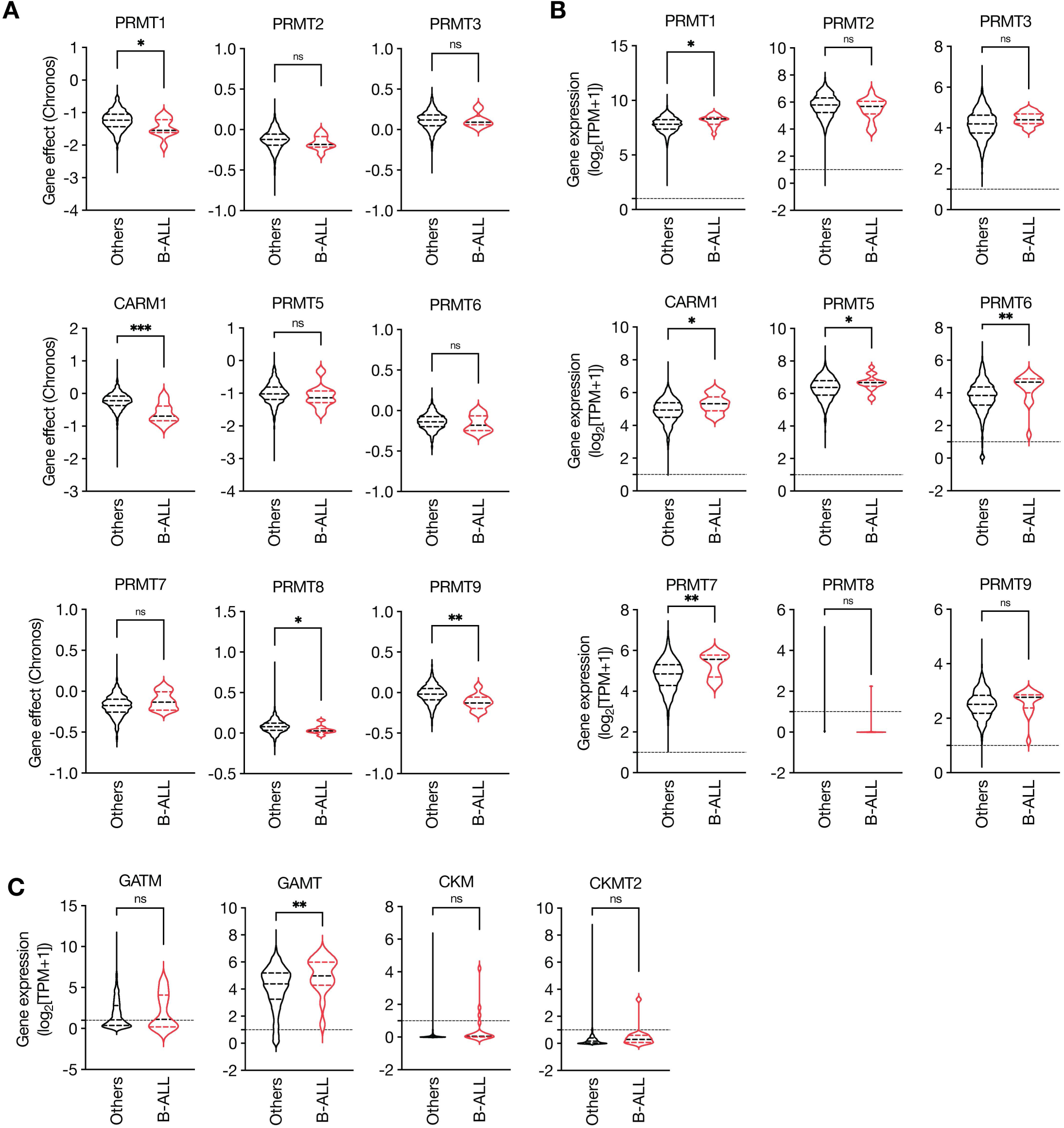
Analysis of gene dependency and expression in B-cell leukemias. **A**, Violin plots of PRMT gene effect distribution from the DepMap 24Q4 release (Chronos). Cell lines were grouped as pediatric B-cell leukemias (n = 10; red) and all other malignancies (n = 1,168; black). **B-C**, Violin plots of gene expression data for PRMT genes (B) or creatine synthesis pathway genes (C) from the DepMap 24Q4 release shown as log_2_ transcripts per million (TPM) +1. Cell lines were grouped as pediatric B-cell leukemias (n = 17; red) and all other malignancies (n = 1,656; black). p-values in were derived from a two-tailed Mann-Whitney test (* p < 0.05; **p < 0.01; ***p < 0.001; ns, not significant).

**Supplementary Figure 5.**
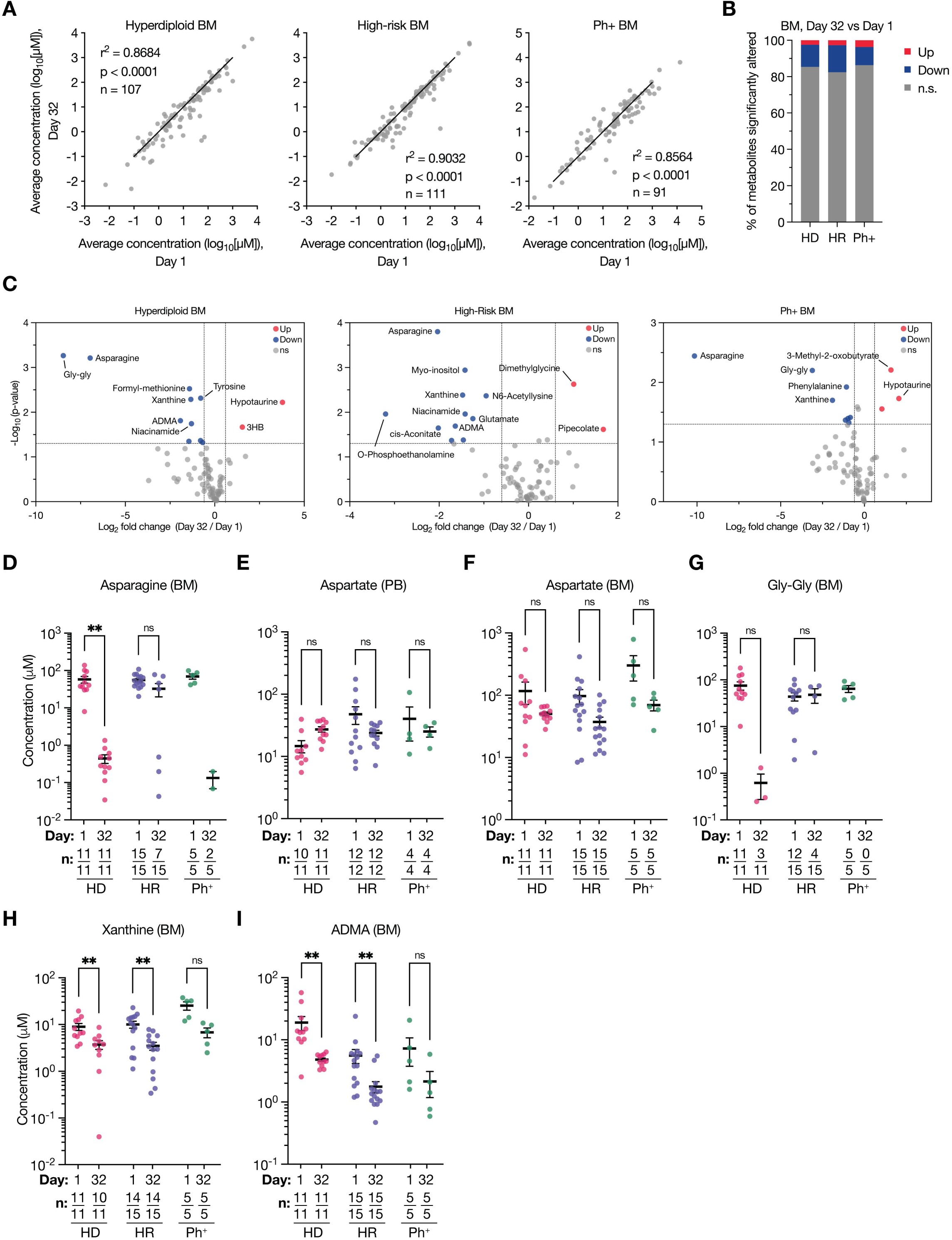
Analysis of metabolite changes in bone marrow plasma of B-ALL patients pre- and post-induction chemotherapy. **A**, Scatter plots of metabolite concentrations measured by LC/MS in bone marrow (BM) plasma from B-ALL patients on day 32 post-induction chemotherapy (day 32) compared to the day of diagnosis (day 1). The number of metabolites measured is indicated in each panel. r and p-values were determined by Pearson correlation. **B**, Bar plot representing the percent of metabolites in bone marrow plasma that were significantly higher (Up) or lower (Down) on day 32 compared to day 1. Cutoffs of | log2 fold change | > 0.6 and raw p-value < 0.05 were used to select significantly altered metabolites. n.s.: not significant. **C**, Volcano plots depicting metabolites that significantly change in concentration in bone marrow plasma between day 1 and day 32 post-induction chemotherapy. Cutoffs of | log_2_ fold change | > 0.6 and raw p-value < 0.05 were used to select significantly altered metabolites. For panels A, C, n = 12 (HD D1), n = 11 (HD D32), n = 19 (HR D1), n = 17 (HR D32), n = 5 (Ph+ D1), n = 5 (Ph+ D32) patient samples. **D-I**, Levels of selected metabolites measured by LC/MS in peripheral blood (PB) or BM plasma from paired B-ALL patients on day 1 or day 32 post-induction chemotherapy. Data are presented as mean ± SEM and p-values were derived from a Wilcoxon matched-pairs signed rank test with Holm-Sidak’s multiple comparisons correction (* p < 0.05; **p < 0.01; ns, not significant). n values indicate the number of samples with detectable metabolite levels (numerator) out of the total number of samples analyzed (denominator). 3HB: 3-hydroxybutyrate; ADMA: asymmetric dimethylarginine; GAA: guanidinoacetic acid; Gly-Gly: glycyl-glycine.

**Supplementary Figure 6.**
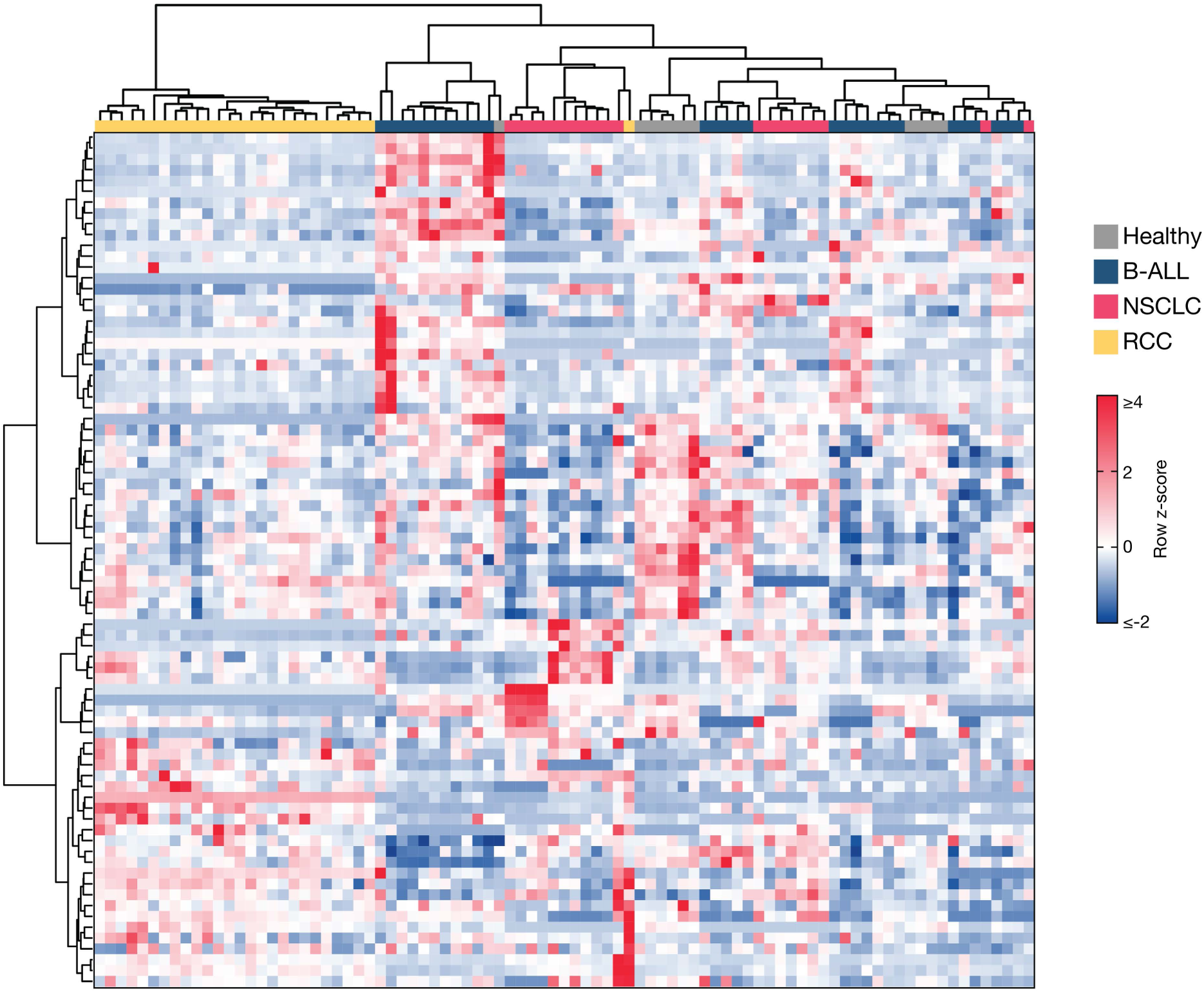
Hierarchical clustering of peripheral blood plasma metabolite composition in B-ALL, NSCLC, and RCC patients. Hierarchical clustering of metabolites measured in peripheral blood plasma from healthy donors and patients bearing B-ALL on day of diagnosis, non-small cell lung cancer (NSCLC), or renal cell carcinoma (RCC). The B-ALL subtypes were collapsed together. Data represent n = 11 (healthy), n = 29 (B-ALL), n = 20 (NSCLC), or n = 27 (RCC) patient samples. Data presented within each column of the heatmap were z-score normalized.

**Supplementary Figure 7.**
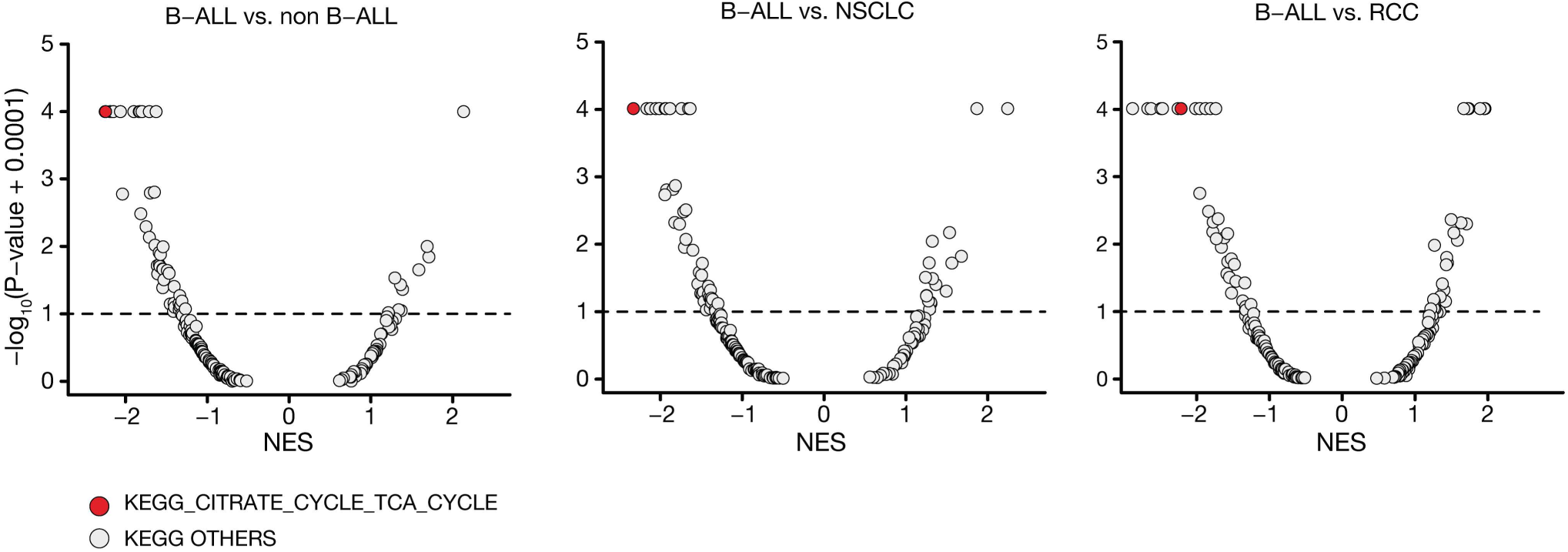
TCA cycle pathway is a dependency in B-ALL. Volcano plots showing enrichment of KEGG pathways for 1,150 cell lines in the Depmap Chronos 24Q2 dataset. The KEGG TCA cycle pathway (red dot) scored as among most depleted in the B-lymphoblastic leukemia/lymphoma lineage (n = 13) compared to all other cell lines (n = 1,137) (left), compared to non-small cell lung cancer (NSCLC) (n = 98) (middle) and renal cell carcinoma (RCC) (n = 29) (right). Normalized enrichment score (NES) shown on x-axis.

